# The nuclear membrane protein Samp1 links peripheral genome organization to the myogenic transcriptional program

**DOI:** 10.64898/2026.09.24.754269

**Authors:** Urška Kašnik, Irene Mei, Silvia Gómez Alcalde, Saira Aftab, Eva Hedlund, Einar Hallberg

## Abstract

Samp1 is an inner nuclear membrane protein required for myogenic differentiation and involved in chromatin organization at the nuclear periphery. Here, we investigated whether these functions are connected by studying the effects of Samp1 depletion during C2C12 myogenic differentiation using immunofluorescence microscopy, RNA sequencing, FRIC, and chromosome-positioning analysis. Samp1-depleted cells showed strongly reduced MyHC expression and virtually abrogated multinucleated fiber formation. Although cell-cycle withdrawal was not prevented, the transcriptional program driving differentiation was drastically perturbed, with reduced muscle-associated transcripts and incomplete repression of genes normally downregulated during myogenesis. Samp1 depletion also disrupted peripheral chromatin organization and prevented the accumulation of peripheral heterochromatin typically seen during differentiation. In addition, radial chromosome distribution was disrupted, evidenced by the failure of chromosome 8 to reposition to the nuclear periphery during differentiation. Together, these findings link the requirement for Samp1 in myogenic differentiation to its role in genome organization at the nuclear periphery.

## Introduction

The nuclear envelope (NE) forms the boundary between the nucleus and the cytoplasm, but its functions extend far beyond compartmentalization and mechanical support (Wilson and Berk, 2010). Proteins of the inner nuclear membrane (INM) and the nuclear lamina interact with chromatin and chromatin-associated proteins and contribute to genome organization and gene regulation at the nuclear periphery (Korfali et al., 2010; Solovei et al., 2013). Proteomic studies have identified a diverse set of several hundred nuclear envelope transmembrane proteins (NETs) (Korfali et al., 2012, 2010; Schirmer et al., 2003; Wilkie et al., 2011). Their expression varies remarkably between tissues, with most NETs showing tissue-specific expression patterns (Korfali et al., 2012). In skeletal muscle, the expression of several NETs increases during myogenic differentiation, suggesting that NE protein composition changes as myoblasts acquire a differentiated state (Wilkie et al., 2011). These findings suggest that NE protein composition is adapted to cell type and can change during differentiation, potentially contributing to cell-type-specific cellular functions.

Skeletal myogenesis is a multistep process in which proliferating myoblasts withdraw from the cell cycle, activate muscle-specific gene expression and ultimately fuse to form multinucleated myotubes. These events are temporally separable: in C2C12 myoblasts, myogenin induction precedes p21-associated cell-cycle withdrawal, followed by expression of myosin heavy chain (MyHC) and subsequent cell fusion (Andrés and Walsh, 1996). Myogenesis is regulated by coordinated transcriptional networks involving MyoD and myogenin, acting together with MEF2 proteins to activate overlapping sets of muscle-specific genes (Blais et al., 2005). RNA-seq analyses further show broad induction of genes involved in muscle development and contractile function together with repression of cell-cycle-associated genes during C2C12 differentiation (Doynova et al., 2017).

These changes in gene expression take place within the spatially organized nucleus. Within the interphase nucleus, transcriptionally active euchromatin is generally enriched in the nuclear interior, whereas the more compact and transcriptionally repressed heterochromatin is enriched at the nuclear periphery and around nucleoli (Solovei et al., 2013). At the nuclear periphery, large genomic regions interact with NE proteins in domains referred to as lamina-associated domains (LADs). Genome-wide mapping of lamina interactions showed that LADs generally have low transcriptional activity and are associated with repressive chromatin marks (Pickersgill et al., 2006; Guelen et al., 2008). Peripheral heterochromatin is maintained through interactions with NE components; for example, LBR- and lamin A/C-dependent mechanisms act at different stages of differentiation to tether heterochromatin to the nuclear periphery (Solovei et al., 2013). Genome–lamina interactions are dynamic and undergo extensive reorganization during differentiation (Peric-Hupkes et al., 2010).

At a larger scale, individual chromosomes occupy discrete territories within the interphase nucleus and show preferred radial positions that can differ between tissues (Parada et al., 2004). These spatial arrangements are not fixed but can change during differentiation. In C2C12 cells, Hi-C analysis showed that the overall genome architecture remains largely conserved during myoblast-to-myotube differentiation, while specific genomic regions undergo changes in chromatin interactions and their association with active or inactive chromatin compartments (Doynova et al., 2017). Reorganization is also evident at the nuclear periphery. For example, chromosome 8 relocates toward the nuclear periphery during C2C12 differentiation, while numerous genomic regions gain or lose association with the nuclear lamina (Robson et al., 2016). NETs can contribute to these spatial changes: different NETs have been shown to influence the positioning of distinct chromosomes (Zuleger et al., 2013), while certain muscle-specific NETs can promote peripheral repositioning of selected genes and contribute to their transcriptional repression during myogenesis (Robson et al., 2016).

Samp1 (Buch et al., 2009), encoded by *TMEM201*, also known as NET5, has been linked to several aspects of spatial genome organization. Samp1 is an INM protein functionally associated with the LINC complex and the A-type lamina network (Gudise et al., 2011). Overexpression of Samp1 was shown to promote the peripheral localization of chromosome 5 in human fibrosarcoma cells (Zuleger et al., 2013). Fluorescence Ratiometric Imaging of Chromatin (FRIC) analysis (Bergqvist et al., 2019) further showed that Samp1 promotes peripheral heterochromatin: depletion of Samp1 reduced heterochromatin at the nuclear periphery in U2OS cells, whereas Samp1 overexpression increased it. Samp1 expression is relatively high in skeletal and cardiac muscle (Korfali et al., 2012; Wilkie et al., 2011), and its protein levels increase approximately sevenfold during C2C12 differentiation (Jafferali et al., 2017). Depletion of Samp1 blocks myotube formation and strongly reduces expression of myogenic markers, while differentiation can be restored by expression of RNAi-resistant human Samp1 (Jafferali et al., 2017). A broader role of Samp1 in differentiation is supported by studies in human induced pluripotent stem cells, where ectopic Samp1 expression induced rapid differentiation even under pluripotency-maintaining conditions (Bergqvist et al., 2017). Together, these studies link Samp1 to both peripheral genome organization and cell differentiation, but whether its effects on these processes are connected remains unclear.

Mutations in genes encoding lamins and other nuclear-envelope proteins cause a diverse group of disorders collectively referred to as laminopathies or nuclear envelopathies, including diseases affecting skeletal and cardiac muscle (Shin and Worman, 2022). Emery-Dreifuss muscular dystrophy (EDMD) is an inherited muscle disorder characterized by early joint contractures, progressive muscle weakness and wasting, and cardiac involvement (Heller et al., 2020). Samp1 directly interacts with the INM protein emerin (Jafferali et al., 2014), a LEM-domain protein with roles in chromatin tethering, gene regulation and mechanotransduction (Berk et al., 2013), and is functionally associated with A-type lamins (Gudise et al., 2011). Mutations in EMD and LMNA, encoding emerin and A-type lamins, respectively, are established causes of EDMD (Muchir and Worman, 2019). In HeLa cells, depletion of Samp1 causes detachment of the centrosome from the NE (Gudise et al., 2011), a phenotype also observed in emerin-deficient EDMD patient fibroblasts and in cells lacking emerin or lamin A/C (Salpingidou et al., 2007). The relevance of the Samp1–emerin interaction to muscle differentiation is further supported by recent work showing that emerin is required for recruitment of centrosomal proteins and proper organization of SUN1 and SUN2 at the NE during human muscle differentiation, processes that are disrupted in emerin-deficient EDMD muscle cells (Mattioli et al., 2026). An additional connection to EDMD comes from the finding that Samp1 localization is disrupted in myotubes carrying disease-associated *LMNA* mutations (Mattioli et al., 2018). Furthermore, three missense variants in *TMEM201*, encoding Samp1 (p.G15A, p.G18S and p.G597S), were identified in patients with EDMD-like phenotypes (Meinke et al., 2020). Although these variants occurred together with variants in other genes, the authors suggested that a *TMEM201* variant could contribute to disease severity in at least one patient. *TMEM201* expression is also reduced in myotonic dystrophy type 1 muscle and correlates with muscle strength, with lower expression associated with greater muscle weakness (Todorow et al., 2023). Together, these findings raise the possibility that Samp1 functions involved in myogenic differentiation and genome organization may also be relevant to muscle disease.

Here, we investigate how Samp1 depletion affects myogenic differentiation and whether these changes are associated with altered spatial genome organization. Using the C2C12 Samp1-knockdown model, we combine analysis of myogenic differentiation with RNA sequencing, FRIC and three-dimensional chromosome 8 DNA-FISH. We show that Samp1 depletion strongly impairs progression through myogenic differentiation and alters the accompanying transcriptional changes. Importantly, Samp1-depleted cells fail to show the increase in peripheral heterochromatin and repositioning of chromosome 8 toward the nuclear periphery observed in control cells during differentiation. Together, our findings support a model in which Samp1 contributes to genome repositioning at the nuclear periphery during myogenic differentiation and thereby influences the expression of genes that normally become repressed during this process.

## Results

### Samp1 depletion impairs myogenic differentiation of C2C12 cells

We previously showed that Samp1 expression increased sevenfold during myogenic differentiation of C2C12 cells and that Samp1 depletion blocked myogenic differentiation (Jafferali et al., 2017). To investigate the mechanisms underlying this differentiation defect, we compared C2C12 cell lines stably expressing an shRNA targeting all Samp1 isoforms (Samp1 KD) with non-targeting control shRNA (Fig. 1a,b). Myogenic differentiation was followed for three days and evaluated by immunofluorescence microscopy, western blotting, and RNA sequencing.

**Figure 1.**
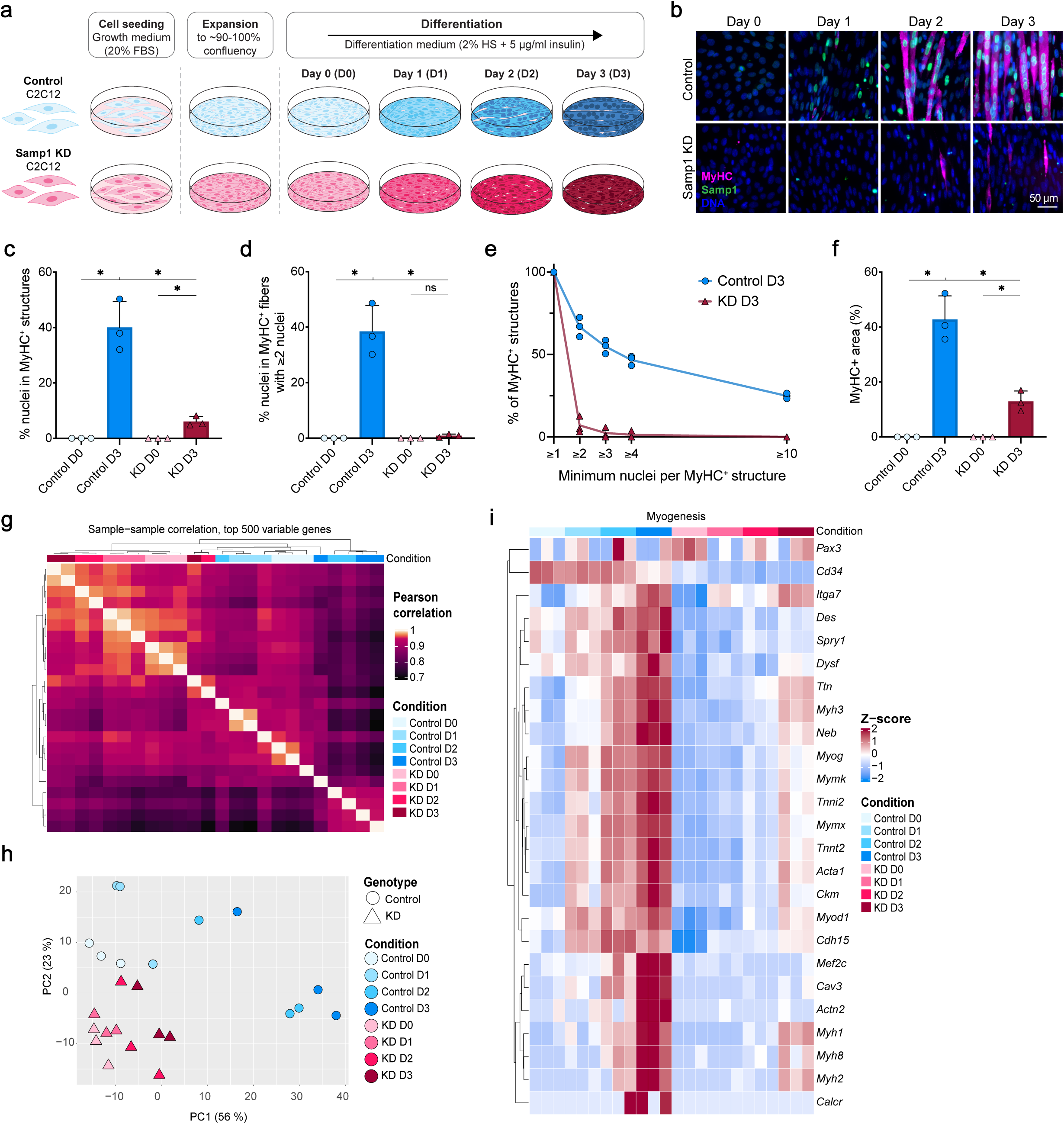
Samp1 depletion impairs myogenic differentiation. Control and Samp1-knockdown (KD) C2C12 cells were differentiated for 3 days. **(a)** Experimental overview. Cells were seeded in growth medium containing 20% fetal bovine serum, expanded to approximately 90–100% confluence, and transferred to differentiation medium containing 2% horse serum and 5 µg/mL insulin at day 0. **(b)** Representative images at days 0–3 showing MyHC (magenta), Samp1 (green), and Hoechst-stained DNA (blue). Scale bar, 50 µm. **(c)** Myogenic index, defined as the percentage of nuclei within MyHC-positive structures. **(d)** Fusion index, defined as the percentage of nuclei within MyHC-positive fibers containing at least two nuclei. **(e)** Percentage of MyHC-positive structures containing at least the indicated number of nuclei at day 3. **(f)** MyHC-positive area as a percentage of total image area. Quantifications in **c–f** were obtained from the MyHC/Ki67/Hoechst dataset shown in Fig. 2b. Three independent biological experiments were analyzed; for **c–e**, at least 1,700 nuclei and 400 MyHC-positive objects were evaluated per condition in each experiment. Data are mean ± SD; symbols represent biological replicates. **(g)** Sample-to-sample Pearson correlation heatmap based on VST-normalized counts, using distance = 1 − Pearson correlation and complete linkage clustering. **(h)** PCA of the 500 most variable genes after z-score normalization. **(i)** Heatmap of row-scaled z-scores of VST-normalized counts for curated myogenic marker genes; columns are ordered by condition (control and Samp1-KD, D0–D3) and rows are hierarchically clustered. RNA-seq analyses in **g–i** used three independent biological replicates per condition and time point. Paired two-tailed t-tests were used for **c, d,** and **f**. Exact P values: **c**, control D0 versus control D3, *P* = 0.0176; KD D0 versus KD D3, *P* = 0.0277; control D3 versus KD D3, *P* = 0.0279; **d**, control D0 versus control D3, *P* = 0.0191; KD D0 versus KD D3, *P* = 0.1123; control D3 versus KD D3, *P* = 0.0177; **f**, control D0 versus control D3, *P* = 0.0130; KD D0 versus KD D3, *P* = 0.0270; control D3 versus KD D3, *P* = 0.0272. \**P* < 0.05; ns, not significant.

Immunofluorescence analysis confirmed efficient Samp1 depletion and revealed a pronounced defect in myogenic differentiation. During differentiation, control cells progressively formed elongated, MyHC-positive multinucleated fibers, whereas Samp1 KD cultures contained only a small number of MyHC-positive cells and showed very limited multinucleation (Fig. 1b). The myogenic index, defined as the percentage of nuclei within MyHC-positive structures, was fivefold lower in KD cells than in control cells (Fig. 1c).

The fusion index, defined as the percentage of nuclei within MyHC-positive fibers containing at least two nuclei, increased significantly during differentiation in control cells but not in Samp1 KD cells (Fig. 1d). At day three, less than 1% of nuclei in KD cultures were located within multinucleated MyHC-positive fibers (Fig. 1d), and very few MyHC-positive structures contained multiple nuclei (Fig. 1e). The total MyHC-positive area was also markedly lower in Samp1 KD than in control cultures (Fig. 1f). Western blotting independently confirmed reduced Samp1 protein abundance in KD cells at both day 0 and day 3. The MyHC protein level was strongly induced during differentiation in control cells but remained low in KD cells, consistent with the immunofluorescence phenotype (Figure 1–figure supplement 1).

To determine whether the impaired differentiation phenotype was accompanied by changes at the transcriptional level, we performed RNA-seq analysis of control and KD cells collected before differentiation and on each day through day three. Pearson correlation analysis showed high consistency among biological replicates, with sample relationships reflecting both differentiation stage and genotype (Fig. 1g). Principal component analysis similarly separated late-stage control samples from undifferentiated and early differentiating samples, whereas KD samples remained closer to undifferentiated control cells (Fig. 1h). Consistent with this pattern, control cells progressively induced the myogenic regulator *Myog*, the fusion-associated genes *Mymk* and *Mymx*, and sarcomeric and contractile genes including *Ttn, Neb, Acta1, Myh1* and *Myh2*, whereas induction of these genes was much weaker in KD cells (Fig. 1i). Together, these results show that Samp1-depleted cells can initiate myogenic differentiation to a limited extent but fail to progress efficiently, resulting in defective MyHC induction, myoblast fusion, and multinucleated fiber formation.

### Samp1 depletion disrupts the myogenic transcriptional program

To define the transcriptional processes affected by Samp1 depletion, we examined coordinated changes in gene modules associated with myogenesis, cell-cycle regulation, extracellular matrix (ECM) organization, signaling, and nuclear-envelope (NE) function. Control cells showed a progressive increase in the myogenic differentiation module from day 0 to day three, accompanied by coordinated changes in cell-cycle and cell-cycle-exit programs (Fig. 2a). These differentiation-associated changes were reduced or delayed in Samp1 KD cells, consistent with their incomplete progression through myogenic differentiation. Analysis of selected cell-cycle- and cell-cycle-exit-associated genes indicated coordinated remodeling of cell-cycle regulation during differentiation in control cells, whereas these changes appeared less coordinated in Samp1 KD cells, particularly at day three (Fig. 2a,e). To determine whether these differences were accompanied by continued proliferation, we quantified Ki67, a nuclear protein widely used as a marker of proliferating cells, at day three (Gerdes et al., 1984). The mean proportion of Ki67-positive nuclei did not differ significantly between control and KD cultures (Fig. 2c). In the combined Ki67/MyHC analysis, Samp1 KD cultures showed a higher mean fraction of non-proliferating cells that had not acquired MyHC expression than control cultures, although this difference did not reach statistical significance (Fig. 2d). As expected from the impaired differentiation phenotype, the MyHC-positive fraction was significantly reduced in Samp1 KD cultures. Together, these results indicate that Samp1 depletion does not simply maintain cells in a proliferative myoblast state but is associated with altered coordination of cell-cycle-associated and myogenic transcriptional programs.

**Figure 2.**
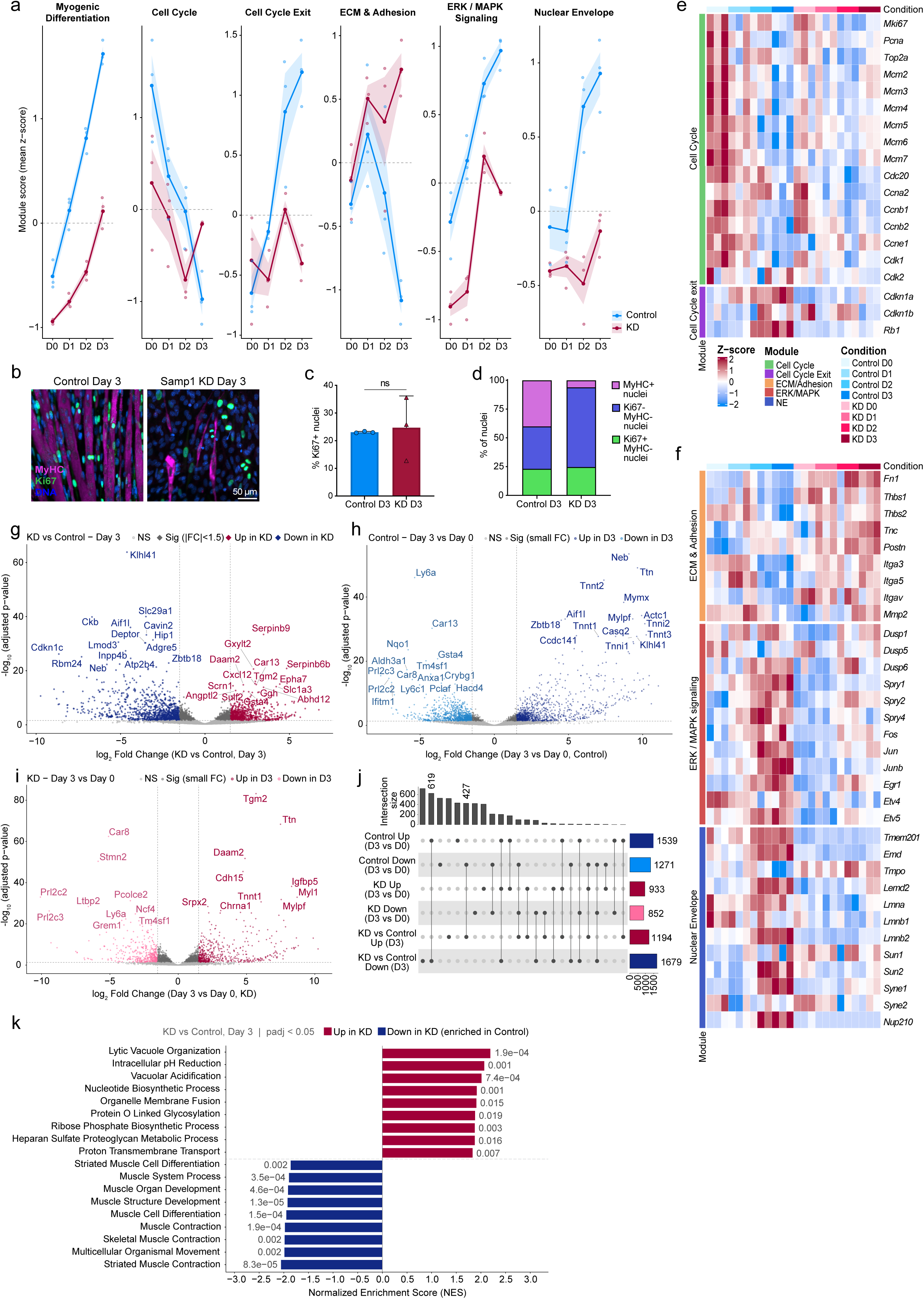
Samp1 depletion disrupts the myogenic transcriptional program. RNA sequencing was performed on control and Samp1-knockdown (KD) C2C12 cells collected at days 0–3 of differentiation. RNA-seq analyses used three independent biological replicates per condition and time point. **(a)** Module scores for myogenic differentiation, cell cycle, cell-cycle exit, extracellular matrix (ECM)/adhesion, ERK/MAPK signaling, and nuclear-envelope functions. Scores are mean z-scores of VST-normalized counts across genes in each module; lines and shading show mean ± SEM and points represent biological replicates. **(b)** Representative day 3 images showing MyHC (magenta), Ki67 (green), and Hoechst-stained DNA (blue). Scale bar, 50 µm. **(c)** Percentage of Ki67-positive nuclei at day 3. **(d)** Day 3 distribution of Ki67-positive/MyHC-negative, Ki67-negative/MyHC-negative, and MyHC-positive nuclei. Three independent biological experiments were analyzed for **c** and **d**, with at least 1,700 nuclei per condition in each experiment. Data in **c** are mean ± SD with biological replicates shown; bars in **d** show mean proportions. Paired two-tailed t-tests showed no difference in the Ki67-positive population (P = 0.8229) or Ki67-negative/MyHC-negative population (P = 0.1210), whereas the MyHC-positive population was reduced in KD cells (P = 0.0303). **(e,f)** Heatmaps of row-scaled z-scores of VST-normalized counts for selected cell-cycle/cell-cycle-exit genes **(e)** and ECM/adhesion, ERK/MAPK, and nuclear-envelope genes **(f)**; columns are ordered by condition without clustering. **(g–i)** Volcano plots showing differential expression between KD and control at day 3 **(g)**, D3 and D0 in control **(h)**, and D3 and D0 in KD **(i)**. DESeq2 Wald tests were used. Genes with |log₂FC| > 1.5 and padj < 0.05 are colored by direction (**g**, red: up in KD; blue: down in KD; **h**, dark blue: up at D3; light blue: down at D3; **i**, dark red: up at D3; pink: down at D3). **(j)** UpSet plot showing overlaps among directional DEG sets using padj < 0.05 and |log₂FC| > 1. **(k)** fgsea results for GO Biological Process terms from the KD versus control day 3 comparison, ranked by normalized enrichment score (NES); positive NES indicates enrichment in KD and negative NES enrichment in control. Only terms with padj < 0.05 are shown; adjusted p-values are indicated.

Samp1 depletion was associated with altered expression of genes involved in ECM organization and adhesion, ERK/MAPK signaling, and nuclear-envelope function (Fig. 2a,f). In control cells, expression of ECM and adhesion genes increased early during differentiation and then declined markedly by day three. In Samp1 KD cells, this decline did not occur and expression remained elevated. This pattern involved ECM components and receptors including *Fn1, Thbs1, Thbs2, Tnc, Postn, Itga3, Itga5* and *Itgav*, together with the matrix-remodeling enzyme *Mmp2*. Genes linked to ERK/MAPK signaling also followed different expression patterns in the two genotypes. In control cells, expression of this group increased progressively during differentiation, whereas the increase was smaller in Samp1 KD cells and did not continue through day 3. Differences were evident in feedback regulators of ERK signaling, including *Dusp1, Dusp5, Duspc, Spry1, Spry2* and *Spry4*, as well as ERK-responsive transcription factors such as *Fos, Jun, Junb, Egr1, Etv4* and *Etv5*. The NE module showed a similarly clear difference between the genotypes. In control cells, the module score increased strongly as differentiation progressed, whereas Samp1 KD cells did not show the same increase. Importantly, this difference extended beyond *Tmem201* (Samp1) itself. Several INM and lamina-associated genes, including *Emd, Lemd2* and *Lmna*, as well as the LINC-complex genes *Sun1, Sun2, Syne1* and *Syne2*, showed higher expression at later stages of control differentiation, whereas the increases were smaller in Samp1 KD cells. *Nup210* also showed a pronounced increase during control differentiation that was much less evident in KD cells. Other genes encoding NE proteins, including *Tmpo, Lmnb1* and *Lmnb2*, showed different expression patterns over the time course (Fig. 2a,f).

Differential expression analysis identified extensive transcriptional differences between control and Samp1 KD cells at day three (Fig. 2g), including reduced expression of several muscle-associated genes in KD cells. At day 3, 1,913 genes were differentially expressed between Samp1 KD and control cells, including 760 genes expressed at higher levels and 1,153 genes expressed at lower levels in KD cells. Comparison of day 3 with day 0 showed substantial transcriptional changes in both genotypes, but fewer genes changed significantly in Samp1 KD cells and the sets of differentially expressed genes only partially overlapped (Fig. 2h–j). In control cells, 2,199 genes changed significantly between days 0 and 3, including 1,206 upregulated and 993 downregulated genes, compared with 1,002 genes in Samp1 KD cells, including 550 upregulated and 452 downregulated genes. The overlap analysis further showed that 619 genes induced during control differentiation were expressed at significantly lower levels in KD than in control cells at day 3. Conversely, 427 genes repressed during control differentiation were expressed at significantly higher levels in KD cells at day 3. Thus, Samp1 depletion altered both the genes normally induced and those normally repressed during myogenic differentiation.

Pathway-enrichment analysis of the day 3 comparison showed that pathways related to muscle-cell differentiation, muscle development and contraction were enriched in control cells, whereas pathways enriched in KD cells included vacuolar acidification and organization, proton transport, nucleotide biosynthesis, glycosylation, and proteoglycan metabolism (Fig. 2k). Together, these analyses show extensive changes in gene expression in Samp1 KD cells after differentiation induction, but the overall pattern differs markedly from that in control cells.

### Samp1 depletion disrupts peripheral chromatin reorganization and chromosome 8 repositioning during myogenic differentiation

Given the previously described role of Samp1 in peripheral genome organization (Bergqvist et al., 2019; Zuleger et al., 2013), we investigated whether its depletion alters spatial chromatin organization during C2C12 differentiation.

Peripheral chromatin organization was examined using FRIC (Bergqvist et al., 2019), based on the ratio of normalized H2B-mCherry to H3.3-EGFP fluorescence (Fig. 3a,b). Higher H2B/H3.3 ratios indicate relatively greater heterochromatin enrichment. In control cells, differentiation produced a clear redistribution of the H2B/H3.3 ratio across the nuclear radius, with increased heterochromatin enrichment in the outermost nuclear zones at day 3 (Fig. 3c). Accordingly, the peripheral heterochromatin index, calculated as the mean H2B/H3.3 ratio in the six outermost zones divided by the mean ratio in the remaining 34 zones, increased between days 0 and 3 in control cells (Fig. 3d).

**Figure 3.**
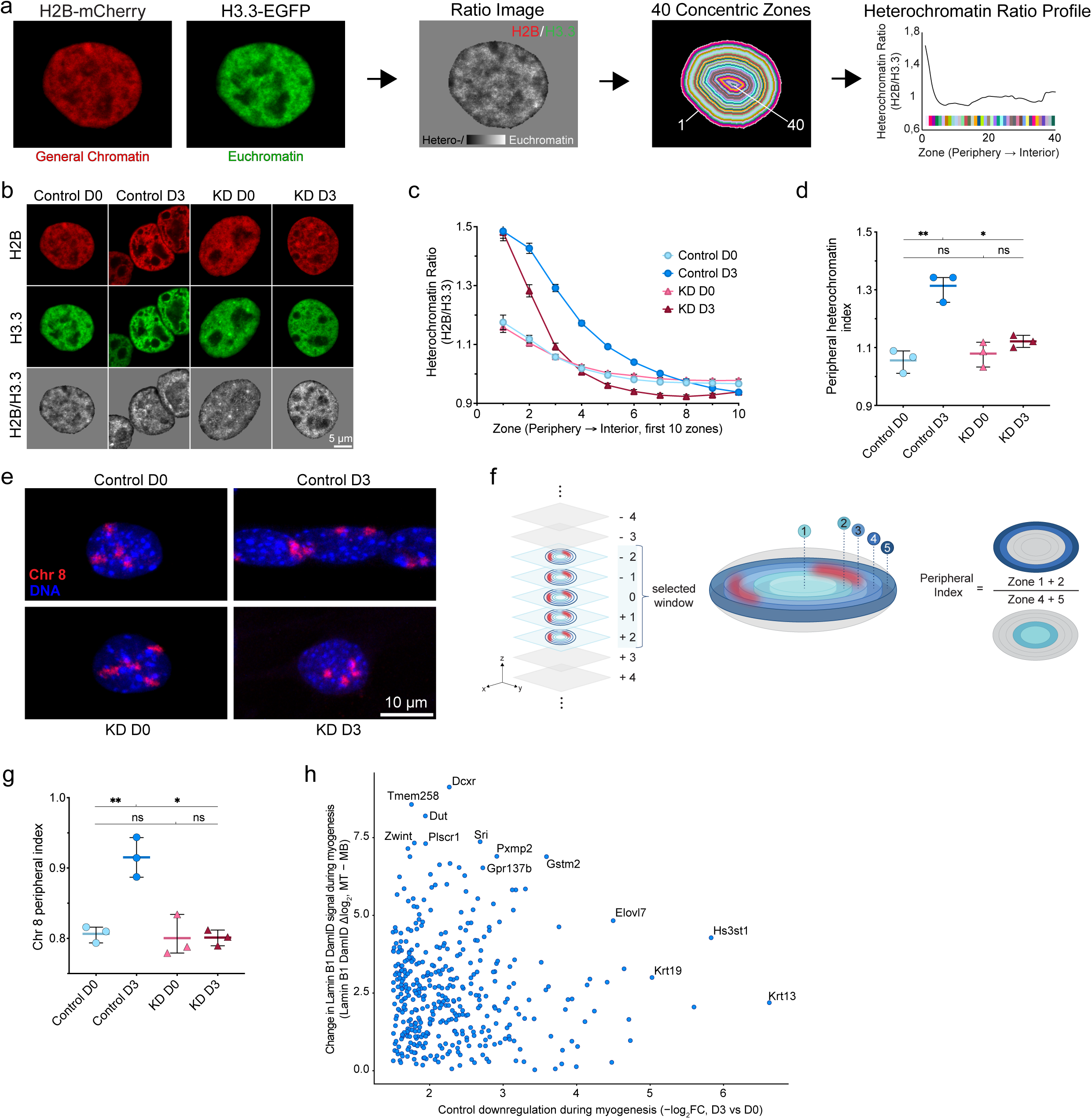
Samp1 depletion disrupts peripheral chromatin reorganization and chromosome 8 repositioning during myogenic differentiation. **(a)** FRIC overview. Normalized H2B-mCherry and H3.3-EGFP images were used to generate H2B/H3.3 ratio images; nuclei were divided into 40 concentric zones from periphery to interior. **(b)** Representative H2B-mCherry, H3.3-EGFP, and ratio images of control and Samp1-knockdown (KD) cells at D0 and D3. Scale bar, 5 µm. **(c)** H2B/H3.3 profiles across the ten outermost zones; lines show mean ± SEM. **(d)** Peripheral heterochromatin index, calculated as the mean ratio in the six outermost zones divided by the remaining 34 zones. Three independent biological experiments were analyzed, with at least 14 nuclei per condition in each experiment. **(e)** Representative chromosome 8 DNA-FISH images showing chromosome 8 (red) and DAPI-stained DNA (blue). Scale bar, 10 µm. **(f)** DNA-FISH workflow. A five-slice window centered on the equatorial section was divided into five concentric equal-volume zones; peripheral index = signal in zones 1 + 2 / signal in zones 4 + 5. **(g)** Chromosome 8 peripheral index. At least 120 nuclei were evaluated per condition in each of three independent biological experiments; control myotube and KD myoblast nuclei were analyzed at day 3. Data in **d** and **g** are mean ± SD with biological replicates shown. **(h)** Scatter plot integrating RNA-seq and published Lamin B1 DamID data (Robson et al., 2016). Points represent genes downregulated in control during differentiation (D3 vs D0, padj < 0.05, log₂FC < −1.5) that did not meet the same criteria in KD and had positive gene-body Δlog₂(myotube − myoblast) Lamin B1 DamID values. Axes show control downregulation (−log₂FC) and change in Lamin B1 DamID signal. Paired two-tailed t-tests were used for **d** and **g**. Exact P values: **d**, control D0 vs D3, P = 0.0012; KD D0 vs D3, P = 0.1679; control vs KD at D0, P = 0.5367; control vs KD at D3, P = 0.0397; **g**, control D0 vs D3, P = 0.0084; KD D0 vs D3, P = 0.9800; control vs KD at D0, P = 0.7086; control vs KD at D3, P = 0.0314. *P < 0.05; **P < 0.01; ns, not significant.

In contrast, Samp1 KD cells did not show a corresponding increase in peripheral heterochromatin. The peripheral heterochromatin index did not change between days 0 and 3 in Samp1 KD cells and was lower than that of control cells at day 3 (Fig. 3c,d). The complete H2B/H3.3 ratio profiles across all 40 nuclear zones are shown in Figure 3–figure supplement 1a.

We next investigated whether Samp1 depletion also affected the radial positioning of an individual chromosome territory. Chromosome 8 was selected because it is rich in genes that are silenced during myogenesis and because previous work identified differentiation-associated repositioning of this chromosome during C2C12 myogenesis (Robson et al., 2016). Chromosome 8 localization was quantified using a three-dimensional DNA-FISH analysis workflow in which fluorescence was measured across five concentric equal-volume nuclear zones (Fig. 3e,f).

In control cells, chromosome 8 redistributed toward the nuclear periphery during differentiation. The peripheral index was calculated as the chromosome 8 signal in the combined peripheral zones 1 + 2 divided by the signal in the combined central zones 4 + 5, with higher values indicating greater peripheral localization. The peripheral index increased significantly between days 0 and 3 (Fig. 3g), as the proportion of chromosome 8 signal in peripheral zones 1 + 2 increased while the proportion in central zones 4 + 5 decreased (Figure 3–figure supplement 1b). No comparable redistribution occurred in Samp1 KD cells: the peripheral index remained unchanged, and the proportion of chromosome 8 signal in the peripheral zones did not increase during differentiation. Accordingly, the peripheral index was lower in Samp1 KD than in control cells at day 3, indicating that chromosome 8 failed to undergo the peripheral repositioning observed during control differentiation.

Finally, we integrated RNA-sequencing data from control cells with published Lamin B1 DamID data from C2C12 myogenesis (Robson et al., 2016). Among genes that were significantly downregulated during differentiation in control cells, but not similarly repressed in Samp1 KD cells, 533 could be matched to the published DamID dataset. Of these, 407 (76.4%) showed higher Lamin B1 DamID signal in myotubes than in myoblasts and were visualized in Fig. 3h. Thus, the majority of matched genes whose repression is impaired in Samp1 KD cells normally become more associated with the nuclear periphery during myogenesis.

## Discussion

In this study, we extend our previous finding that Samp1 is required for C2C12 myogenic differentiation (Jafferali et al., 2017) by showing that the differentiation defect is accompanied by broad changes in gene expression and a failure to reorganize the genome at the nuclear periphery. Samp1-depleted myoblasts initiated MyHC expression to a limited extent but almost completely failed to fuse into multinucleated fibers. Although Samp1-depleted cells responded to differentiation conditions, the differences in gene expression compared to control cells became greater as differentiation progressed. Samp1-depleted cells also failed to increase peripheral heterochromatin or reposition chromosome 8 toward the nuclear periphery during differentiation. In parallel, many genes repressed during differentiation of control cells were not similarly repressed in Samp1-depleted cells. Many of these genes normally increase their association with the nuclear periphery during myogenesis. Together, these findings connect the requirement for Samp1 during myogenic differentiation with the reorganization of the genome at the nuclear periphery.

Cell-cycle withdrawal, muscle-specific gene expression and fusion are temporally separable events during C2C12 differentiation (Andrés and Walsh, 1996), and our results suggest that Samp1 depletion affects progression across several of these steps. In our previous study (Jafferali et al., 2017), Samp1-depleted C2C12 cells showed increased 5-bromo-2′-deoxyuridine (BrdU) incorporation during early differentiation, consistent with delayed cell-cycle withdrawal. By day three, however, the similar mean Ki67-positive fractions in the two genotypes indicate that continued proliferation cannot account for the severe differentiation defect. Together with the altered cell-cycle and cell-cycle-exit transcriptional programs observed here, these findings suggest that Samp1 depletion disrupts the normal coordination between cell-cycle regulation and acquisition of the myogenic program, rather than simply maintaining cells in a proliferative myoblast state.

The RNA-seq data further show that Samp1-depleted cells did not simply remain in their starting myoblast state. Gene expression changed substantially after differentiation induction, but both the normal activation and repression of genes during myogenesis were altered. In our data, control cells strongly induced myogenic regulators, fusion-associated genes and genes involved in sarcomeric and contractile functions, whereas induction of these genes was much weaker in Samp1-depleted cells. This pattern is consistent with previous transcriptomic analysis of normal C2C12 differentiation, which similarly showed strong induction of *Myog, Acta1* and several myosin and troponin genes, together with repression of cell-cycle genes (Doynova et al., 2017). The pathway analysis showed a similar pattern. Control cells were enriched for pathways related to muscle differentiation, muscle development and contraction. In Samp1-depleted cells, the enriched pathways instead included vacuolar acidification, nucleotide biosynthesis and glycosylation. These pathways did not suggest a coherent alternative transcriptional program but showed that Samp1-depleted cells respond to differentiation conditions without following the normal course of myogenesis. Their response is therefore altered rather than simply delayed.

The altered expression of ECM and adhesion genes may be particularly relevant to the failure of myoblast fusion. Dynamic regulation of cell–matrix interactions is important during myogenesis. C2C12 myoblasts normally show transient changes in adhesion and migration before fusion (Glenn et al., 2010), and fibronectin promotes directional migration and alignment that facilitates myoblast fusion (Vaz et al., 2012). This is particularly relevant to Samp1 because it is involved in nuclear movement during fibroblast polarization and migration, where it associates with SUN2 and lamin A/C (Borrego-Pinto et al., 2012). Time-lapse microscopy in our previous study also showed a somewhat slower migration in Samp1-depleted C2C12 cells than in control cells, although both populations remained mobile throughout the observation period (Jafferali et al., 2017). Persistent expression of ECM and adhesion genes in Samp1-depleted cells may therefore reflect incomplete remodeling of cell–matrix interactions during differentiation and could contribute to the failure of myoblast fusion.

The altered expression of genes linked to ERK/MAPK signaling is consistent with our previous finding that Samp1-depleted cells show increased ERK phosphorylation shortly after differentiation induction (Jafferali et al., 2017). A similar connection between NE proteins and ERK signaling during myogenesis has been described for LEMD2/NET25 and emerin. NET25 depletion causes transient ERK1/2 hyperactivation at the start of C2C12 differentiation, and brief inhibition of ERK signaling rescues myogenesis in NET25- and emerin-depleted cells (Huber et al., 2009). Together, these findings suggest that ERK signaling is not properly regulated during differentiation and may contribute to the differentiation defect in Samp1-depleted cells.

The altered expression of genes encoding NE proteins may be functionally important because changes in NE composition can influence myogenic differentiation. A broader difference in the expression of genes encoding NE proteins is also evident in our data: the NE module increased strongly during control differentiation, whereas the increase was much smaller in Samp1-depleted cells. However, these transcriptional changes should not be assumed to reflect equivalent changes in NE protein abundance within the same time period. INM proteins can persist for several days and show considerable variation in turnover (Buchwalter et al., 2019), so changes in mRNA levels during the three-day differentiation period may precede changes in some of the corresponding proteins.

Peripheral genome organization can be functionally important during differentiation; for example, recent work identified LBR and membrane-bound LAP2 as major heterochromatin tethers at the nuclear periphery and showed that their loss disrupts peripheral heterochromatin and impairs differentiation of mouse embryonic stem cells (Lewis et al., 2026). Samp1-depleted cells failed to show the increase in peripheral heterochromatin observed during control differentiation. Previous FRIC analysis showed that Samp1 also influences peripheral chromatin organization in U2OS cells, where Samp1 depletion reduced peripheral heterochromatin enrichment and Samp1 overexpression increased it (Bergqvist et al., 2019). Our results therefore extend the role of Samp1 in peripheral chromatin organization to myogenic differentiation.

The effect of Samp1 depletion was also evident at the level of chromosome positioning. Robson et al. (2016) showed that chromosome 8 normally moves toward the nuclear periphery during C2C12 differentiation. We observed a similar peripheral movement in control cells, whereas chromosome 8 remained unchanged in Samp1-depleted cells. These findings indicate that Samp1 is required for the normal repositioning of chromosome 8 during myogenic differentiation.

Many genes that were repressed during control differentiation but not similarly repressed in Samp1-depleted cells also showed higher Lamin B1 DamID signal in myotubes than in myoblasts in the published data from Robson et al. (2016). Higher Lamin B1 DamID signal indicates increased association with the nuclear periphery, suggesting that these genes normally gain peripheral association during myogenesis. The overlap therefore raises the possibility that the failure to establish normal peripheral genome organization in Samp1-depleted cells contributes to the impaired repression of these genes.

Taken together with the FRIC and chromosome 8 results, these findings strengthen the connection between Samp1 and genome organization at the nuclear periphery. Samp1 depletion prevents spatial changes that normally occur during myogenic differentiation, while the transcriptional response increasingly diverges. Although the present experiments do not establish the molecular mechanism by which Samp1 influences chromatin organization, they show that loss of Samp1 is sufficient to disrupt both genome organization at the nuclear periphery and the normal transcriptional response during myogenic differentiation.

These findings may also be relevant to EDMD, for which the underlying mechanism remains unclear and may involve defects in mechanotransduction, gene regulation or both (Meinke et al., 2020). Several EDMD-associated NE proteins interact with Samp1, including emerin, lamin A and SUN1 (Gudise et al., 2011; Jafferali et al., 2014). In one patient with a very early-onset EDMD-like phenotype, *TMEM201*, encoding Samp1, was mutated (p.G15A) together with a non-causative mutation in *LMNA* (Meinke et al., 2020). These genetic links to EDMD and Samp1 interactions with EDMD-associated NE proteins, together with the strong effects of Samp1 depletion on chromatin organization and the myogenic transcriptional program presented here, support misregulation of muscle genes as a possible mechanism contributing to EDMD pathogenesis.

Together, our findings link the requirement for Samp1 in myogenic differentiation to its role in organizing the genome at the nuclear periphery.

## Materials and methods

### Antibodies

Primary antibodies used for immunofluorescence were rabbit anti-Ki67 (Abcam, Cambridge, UK, ab15580; 1:500), mouse anti-myosin heavy chain (MyHC; MF20, Developmental Studies Hybridoma Bank, Iowa City, IA, AB_2147781; 1:50), and rabbit anti-Samp1a (1:400), described previously (Gudise et al., 2011).

The following fluorophore-conjugated secondary antibodies were used at 1:5,000: Alexa Fluor 488 donkey anti-rabbit IgG (H+L) (Invitrogen, Thermo Fisher Scientific, Waltham, MA, A21206), Alexa Fluor 568 goat anti-mouse IgG (H+L) (Invitrogen, A11004), and Alexa Fluor 647 donkey anti-mouse IgG (H+L) (Invitrogen, A31571). Alexa Fluor 647 donkey anti-mouse antibody was used for MyHC detection in pTandemH-expressing samples.

Primary antibodies used for western blotting were rabbit anti-Samp1a, described previously (Gudise et al., 2011), mouse anti-MyHC (RCD Systems, Bio-Techne, Minneapolis, MN, MAB4470), and mouse anti-β-actin (Thermo Fisher Scientific, MA5-15739). Primary antibodies were diluted 1:200. HRP-conjugated anti-mouse IgG (Cytiva, Marlborough, MA, NA931) and goat anti-rabbit IgG (H+L) (Thermo Fisher Scientific, A16096) were diluted 1:1,000.

### Cell culture and differentiation

Parental C2C12 mouse myoblasts were obtained from the American Type Culture Collection (ATCC, Manassas, VA, USA; CRL-1772; RRID:CVCL_0188). ATCC authenticates C2C12 cells by cytochrome c oxidase I (COI) species determination. Stable C2C12 mouse myoblast lines expressing either a non-targeting control shRNA or an shRNA targeting Samp1 (Samp1 sh2) were generated and characterized previously (Jafferali et al., 2017). The Samp1 sh2 line was previously validated alongside an independent Samp1-targeting shRNA, and the impaired myogenic differentiation phenotype was rescued by expression of RNAi-resistant human Samp1. The Samp1 sh2 sequence, which targets all murine Samp1 isoforms, was 5′-GAGCAGTACAATGGCTTTCAA-3′. These cell lines are referred to throughout as control and Samp1 knockdown (KD), respectively. Cultures were routinely tested for mycoplasma contamination and were negative during the experiments reported here.

Both cell lines were maintained in high-glucose DMEM (Gibco, Thermo Fisher Scientific, Waltham, MA, 41965039) supplemented with 20% FBS (Gibco, A5256801) and 1% penicillin–streptomycin (Gibco, 15140122). Cells were cultured at 37 °C in a humidified atmosphere containing 5% CO₂.

To induce myogenic differentiation, cells were grown to 90–100% confluence, and the growth medium was replaced with differentiation medium consisting of high-glucose DMEM supplemented with 2% heat-inactivated horse serum (Gibco, 26050070), 1% penicillin–streptomycin, and 5 µg/mL insulin (Sigma-Aldrich, St. Louis, MO). Approximately 70% of the medium was replaced with fresh differentiation medium every 24 h. Undifferentiated cultures were collected or fixed immediately before differentiation induction (day 0), whereas differentiated cultures were collected or fixed at the indicated time points up to day 3. For microscopy experiments, cells were grown in 24-well imaging plates (ibidi GmbH, Gräfelfing, Germany, 82426).

Unless otherwise indicated, experiments were performed using three independent biological replicates. A biological replicate comprised an independent experiment performed on a separate day using independently cultured cells.

### Transfection for FRIC experiments

For FRIC experiments, C2C12 cells were transfected in suspension before seeding using the GenJet C2C12 in vitro DNA transfection reagent (SignaGen Laboratories, Frederick, MD, SL100489-C2C12-05) and the previously described bicistronic pTandemH reporter plasmid, which coexpresses H2B-mCherry and H3.3-EGFP (Bergqvist et al., 2019). Transfections were performed using a 4:1 reagent-to-DNA ratio according to the manufacturer’s instructions.

At 24 h after transfection, one set of undifferentiated cultures was fixed, whereas differentiation was initiated in the remaining cultures by replacing growth medium with differentiation medium. Differentiated cultures were fixed after 3 days in differentiation medium using 4% paraformaldehyde for 20 min.

### Immunofluorescence staining and microscopy

Cells were fixed with 4% paraformaldehyde for 15–20 min at 4 °C and permeabilized with 0.5% Triton X-100 in PBS for 8–10 min. Samples were blocked for 1 h at room temperature or overnight at 4 °C in PBS containing 2% BSA and 0.05% Tween-20.

Primary antibodies were diluted in blocking solution and incubated with the samples overnight at 4 °C. After washing with PBS containing 0.05% Tween-20, samples were incubated for 1 h at room temperature with the appropriate Alexa Fluor-conjugated secondary antibodies. For pTandemH-expressing samples stained for MyHC, Alexa Fluor 647 donkey anti-mouse IgG was used. Nuclear DNA was counterstained with Hoechst 33342 (Invitrogen, H3570) at 1 µg/mL, corresponding to a 1:10,000 dilution of the supplied 10 mg/mL stock.

Following washing, samples were mounted using Fluoromount-G mounting medium (Invitrogen, 00-4958-02) and covered with glass coverslips. Widefield fluorescence images were acquired using an Axio Observer 7 microscope (Carl Zeiss Microscopy GmbH, Jena, Germany) equipped with a Plan-Apochromat 40×/0.95 Korr M27 air objective. Confocal images were acquired using a Zeiss LSM 700 microscope equipped with a Plan-Apochromat 63×/1.40 Oil DIC M27 objective.

### Immunofluorescence image analysis

Widefield fluorescence images acquired in the Hoechst, MyHC, and Ki67 channels were analyzed using a custom image-analysis pipeline implemented in Python v3.10.19. Nuclei were segmented from the Hoechst channel using Cellpose v3. Nuclei touching image boundaries, poorly segmented nuclei, and segmentation artifacts were excluded.

MyHC-positive objects were segmented from the MyHC channel using ilastik v1.4.0.post1, and the resulting masks were imported into the analysis pipeline. Nuclei were assigned to MyHC-positive objects based on their centroid position and overlap with the corresponding mask. Automated segmentations, nucleus-to-object assignments, and nuclei counts per object were visually inspected and manually corrected when necessary.

The myogenic index was calculated as the percentage of valid nuclei located within MyHC-positive objects. The fusion index was calculated as the percentage of valid nuclei located within MyHC-positive objects containing at least two nuclei. Multinucleation was determined from the number of nuclei assigned to each MyHC-positive object. For presentation of the biological results, segmented MyHC-positive objects are referred to as MyHC-positive structures, whereas objects containing at least two nuclei are referred to as multinucleated fibers.

For MyHC-area measurements, background was removed from the MyHC channel using rolling-ball subtraction. MyHC-positive pixels were identified using a fixed intensity threshold selected separately for each biological replicate and applied uniformly to all experimental groups within that replicate. MyHC-positive area was expressed as a percentage of the total image area.

Ki67-positive nuclei were classified based on nuclear Ki67 intensity and enrichment relative to the local perinuclear background. Automated classifications were visually reviewed and manually corrected when necessary. The Ki67 index was calculated as the percentage of valid nuclei classified as Ki67 positive. Combined Ki67/MyHC categories were derived from the corresponding nuclear classifications. Measurements were calculated per image and summarized by biological replicate.

### FRIC analysis

Confocal images of cells expressing the pTandemH reporter were acquired in the H2B-mCherry and H3.3-EGFP channels. FRIC analysis was performed using a CellProfiler v4.2.6 image-analysis pipeline adapted from Bergqvist et al. (2019). For each pTandemH-expressing nucleus, the equatorial optical section was analyzed.

Nuclei were segmented, and the H2B-mCherry and H3.3-EGFP channels were normalized within each nucleus to account for differences in mean fluorescence intensity and variance between the two channels. A heterochromatin-ratio image was generated by dividing the normalized H2B-mCherry signal by the normalized H3.3-EGFP signal. In contrast to the H3.3/H2B ratio used in the original FRIC study, the heterochromatin ratio in this study was defined as H2B/H3.3, such that higher values indicated relatively greater heterochromatin enrichment.

Each nucleus was divided into 40 concentric zones of equal width extending from the nuclear periphery to the interior, and the mean H2B/H3.3 ratio was calculated within each zone. The peripheral heterochromatin index was calculated as the mean ratio in the six outermost zones divided by the mean ratio in the remaining 34 inner zones. Measurements were obtained for individual nuclei and summarized by biological replicate.

### Chromosome 8 DNA fluorescence in situ hybridization

C2C12 cells were grown in 24-well imaging plates and fixed either immediately before differentiation induction or after 3 days of differentiation. Cells were fixed with 4% paraformaldehyde for 10–15 min and permeabilized with 0.5% Triton X-100 for 10 min.

Samples were incubated in 0.1 M HCl for 5 min, treated with PureLink RNase A (Invitrogen, 12091021) at 100 µg/mL in PBS for 1 h at 37 °C, and digested with pepsin from porcine stomach mucosa (Sigma-Aldrich, P7000) at 0.05 mg/mL in 0.01 M HCl for 3 min at 37 °C. After pepsin treatment, samples were washed with PBS containing 5 mM MgCl₂.

Samples were dehydrated sequentially in 70%, 90%, and 100% ethanol for 3 min at each concentration and allowed to air-dry. An XMP 8 Orange whole-chromosome paint probe (MetaSystems Probes, Altlussheim, Germany, D-1405-050-OR 10) was applied to each sample and sealed using CytoBond removable coverslip sealant (SciGene, Santa Clara, CA, 2020-00-1). After the sealant had solidified, the probe and cellular DNA were co-denatured by heating the imaging plate to 78 °C for 3 min. Hybridization was performed in a humidified chamber at 37 °C for approximately 48 h.

Following hybridization, samples were washed in 0.4× SSC for 2 min at 72 °C, followed by 2× SSC containing 0.05% Tween-20 for 30 s at room temperature. Samples were rinsed with Milli-Q water, gently air-dried, and mounted using Fluoromount-G mounting medium containing DAPI (Invitrogen, 00-4959-52). Confocal z-stacks were acquired at 0.4-µm intervals.

### DNA-FISH image analysis

Fluorescence z-stacks were analyzed using a custom image-analysis pipeline implemented in Python v3.9.23. Nuclei were segmented from the DAPI channel using Cellpose v3 on individual optical sections, and the resulting two-dimensional masks were linked across consecutive z-slices to generate three-dimensional nuclear objects. Segmentation overlays were inspected manually, and nuclei that touched image boundaries, were poorly segmented, contained segmentation artifacts, or overlapped with neighboring nuclei were excluded.

In differentiated cultures, nuclei were manually classified as myoblast or myotube nuclei according to their cellular context. Control D3 myotube nuclei and KD D3 myoblast nuclei were retained for the final analysis.

For each nucleus, an equatorial optical section was selected automatically from the DAPI channel based on nuclear sharpness, DAPI intensity, and cross-sectional area. This section and two adjacent sections on either side were retained, generating a standardized five-slice analysis volume. Nuclei lacking a complete five-slice window were excluded.

Chromosome 8 fluorescence was corrected for local background by subtracting the 20th-percentile intranuclear intensity separately for each nucleus and optical section, with negative values set to zero. Corrected, non-thresholded fluorescence intensity was quantified within five concentric equal-volume 3D nuclear zones, with zone 1 representing the nuclear periphery and zone 5 the nuclear center. Signal within each zone was expressed as a proportion of the total corrected intranuclear chromosome 8 fluorescence. The peripheral index was calculated as:

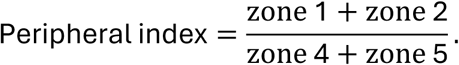

Higher values indicated greater enrichment of chromosome 8 at the nuclear periphery. Measurements were obtained for individual nuclei and summarized by biological replicate.

### Western blotting

Control and Samp1-knockdown (KD) C2C12 cells were collected before differentiation induction and after 3 days of differentiation. Cells cultured in 100-cm² dishes were lysed directly in approximately 500 µL RIPA buffer (Thermo Scientific, Thermo Fisher Scientific, Waltham, MA, 89900). Total protein concentration was determined using a bicinchoninic acid protein assay according to the manufacturer’s instructions and adjusted to 2 mg/mL.

Equal amounts of total protein were separated on 12% SDS–polyacrylamide gels and transferred to 0.2-µm nitrocellulose membranes using Tris–glycine transfer buffer containing 25 mM Tris base, 192 mM glycine, and 20% methanol.

For Samp1 detection, membranes were blocked for 1 h at room temperature in Tris-buffered saline containing 0.1% Tween-20 (TBST) and 5% nonfat dry milk. For MyHC detection, membranes were blocked in TBST containing 3% BSA. Membranes were incubated overnight at 4 °C with rabbit anti-Samp1a antibody, described previously, mouse anti-MyHC antibody (RCD Systems, MAB4470), or mouse anti-β-actin antibody (Thermo Fisher Scientific, MA5-15739). Primary antibodies were diluted 1:200 in TBST containing 2% nonfat dry milk or 2% BSA, as appropriate.

After primary-antibody incubation, membranes were washed three times with TBST and incubated for 1 h at room temperature with HRP-linked anti-mouse IgG (Cytiva, NA931) or goat anti-rabbit IgG (H+L) HRP-conjugated secondary antibody (Thermo Fisher Scientific, A16096), diluted 1:1,000 in TBST containing 2% nonfat dry milk or BSA. Membranes were subsequently washed three times with TBST.

Protein bands were visualized using Pierce ECL Western Blotting Substrate (Thermo Scientific, 32106) and imaged using an Azure 600 Western Blot Imaging System (Azure Biosystems, Dublin, CA). Band intensities were quantified using ImageJ v1.54p. Samp1 and MyHC band intensities were normalized to the corresponding β-actin loading-control intensity. For graphical presentation, Samp1 values were expressed relative to control day 0 and MyHC values relative to control day 3. Normalized biological-replicate values were analyzed using GraphPad Prism version 11. Western blotting was performed using three independent biological replicates for MyHC and four for Samp1.

### Bulk RNA-seq library preparation using Smart-Seq2

Bulk RNA-seq libraries were generated using a modified Smart-Seq2 protocol (Picelli et al., 2014). Cells were lysed in a solution containing 2% Triton X-100 (Sigma-Aldrich, T9284-100 mL), 2 U/µL recombinant RNase inhibitor (Takara Bio Inc., Kusatsu, Shiga, Japan, 2313A), and 1 mM DTT. For each reaction, 5 µL lysate was incubated with 1.63 µL 10 µM SMARTer oligo-dTVN primer (Integrated DNA Technologies, Coralville, IA; 5′-biotin-AAGCAGTGGTATCAACGCAGAGTAC(T)₃₀VN-3′), 0.65 µL 25 mM dNTP mix (Thermo Fisher Scientific, R1121), 0.08 µL 100 mM DTT, and 0.14 µL of a 4 × 10⁻⁶ dilution ERCC RNA Spike-In Control Mix (Thermo Fisher Scientific, 4456740). Samples were denatured at 72 °C for 3 min and immediately chilled on ice.

First-strand cDNA synthesis was performed using SuperScript II reverse transcriptase (Thermo Fisher Scientific, 18064014), with each reaction containing 3.25 µL 5 M betaine (Sigma-Aldrich, B0300-1VL), 0.74 µL 100 mM DTT, 0.1 µL 1 M MgCl₂ (Thermo Fisher Scientific, AM9530G), 0.41 µL recombinant RNase inhibitor, and 0.16 µL 100 µM SMARTer TSO-LNA template-switching oligonucleotide (biomers.net GmbH, Ulm, Germany; 5′-biotin-AAGCAGTGGTATCAACGCAGAGTACrGrG+G-3′). Reverse transcription was performed at 42 °C for 90 min, followed by 10 cycles of 50 °C for 2 min and 42 °C for 2 min, and a final incubation at 70 °C for 15 min.

Second-strand synthesis and full-length cDNA amplification were performed using 2× KAPA HiFi HotStart ReadyMix (Roche Diagnostics, Mannheim, Germany, KK2602) and a custom SMARTer ISPCR primer (Integrated DNA Technologies; 5′-AAGCAGTGGTATCAACGCAGAGT-3′). Amplification consisted of an initial denaturation step at 98 °C for 3 min, followed by 18 cycles of 98 °C for 20 s, 67 °C for 15 s, and 72 °C for 6 min, and a final extension at 72 °C for 5 min. Amplified cDNA was purified using 19.5% PEG-based SPRI beads at a 0.85× bead-to-sample ratio and eluted in 25 mM Tris-HCl (pH 8.0; Thermo Fisher Scientific, AM9849).

Sequencing libraries were prepared from 1 ng of amplified cDNA using a low-volume Nextera tagmentation protocol. For each reaction, cDNA was combined with 2.0 µL of 2× TD buffer and 0.1 µL of Nextera Tagment DNA enzyme. Tagmentation was performed at 55 °C for 7 min. Reactions were immediately placed on ice and inactivated by adding 1 µL of 0.2% SDS, followed by incubation at room temperature for 5 min.

Tagmented cDNA was indexed by adding 1 µL each of 1 µM Nextera i5 and i7 index primers. Index ligation and enrichment PCR were performed using Phusion High-Fidelity DNA polymerase (2 U/µL, Thermo Fisher Scientific, F530L). Library amplification consisted of an initial extension at 72 °C for 3 min and denaturation at 95 °C for 30 s, followed by 12 cycles of 95 °C for 10 s, 55 °C for 30 s, and 72 °C for 30 s, and a final extension at 72 °C for 5 min. Indexed libraries were purified using 24% PEG-based magnetic beads and eluted in 25 mM Tris-HCl (pH 8.0). The concentration of each library was determined using Qubit with the dsDNA high sensitivity kit (Thermo Fisher Scientific, Q33231), and library quality was assessed using an Agilent Bioanalyzer (Agilent Technologies, Santa Clara, CA). The resulting libraries were combined into a single pool and sequenced on one lane of a NovaSeq X 10B 300-cycle flow cell (Illumina, San Diego, CA) at the National Genomics Infrastructure (NGI) Sweden, Stockholm.

### RNA sequencing and bioinformatic analysis

RNA sequencing was performed on three independent biological replicates per condition and time point. RNA-seq reads were mapped to the mm39 mouse genome assembly using STAR (v2.7.0e) (Dobin et al., 2013). Expression levels were quantified using rpkmforgenes.py (https://sandberg.cmb.ki.se/rnaseq) with Ensembl gene annotation. All downstream analyses were performed in R (v4.4.3). For quality control and normalization, genes with at least 10 counts in at least three samples were retained. Variance-stabilizing transformation (VST) was applied blind to experimental condition using the vst() function in DESeq2 (v1.46) (Love et al., 2014). Sample-to-sample Pearson correlation was computed on VST-normalized counts and visualized as a hierarchical clustering heatmap using ComplexHeatmap (v2.22) (Gu et al., 2016), with distance defined as 1 − Pearson correlation and complete linkage clustering. Principal component analysis (PCA) was performed on the 500 most variable genes following z-score normalization.

### Differential expression analysis

Two separate differential expression comparisons were performed using DESeq2. First, to identify genes differentially expressed between day 3 and day 0 within each genotype, we retained genes with at least 5 counts in at least 3 samples. A Wald test was applied using the day 3 versus day 0 contrast separately for control and Samp1-KD samples. Second, to identify genes differentially expressed between Samp1-KD and control cells at day 3, a group model approach was used in which genotype and timepoint were merged into a single variable (group = genotype_time), and KD_d3 vs ctrl_d3 was used as the contrast. Here we retained genes with at least 10 counts in at least 3 samples. For the differential expression analyses, genes were considered differentially expressed at adjusted p-value < 0.05 (Benjamini–Hochberg correction) and |log₂FC| > 1.5. For the UpSet overlap analysis, gene sets were defined using adjusted p-value < 0.05 and |log₂FC| > 1. Gene symbols were mapped from Ensembl IDs using the org.Mm.eg.db Bioconductor annotation package (v3.20).

### Gene set and module score analysis

To investigate correlated transcriptional changes across biological programs, module scores were computed for six gene sets: myogenic differentiation, cell cycle, cell cycle exit, ECM/adhesion, ERK/MAPK signaling, and nuclear envelope. Gene sets were defined based on established marker genes and converted to Ensembl IDs using biomaRt (v2.62.1) (Durinck et al., 2009). The genes included in each module are shown in the corresponding heatmaps (Fig. 1i and Fig. 2e,f). For each module, VST-normalized expression values were z-scored per gene across all samples, and the mean z-score across module genes was computed per sample. Module scores were summarized across timepoints (d0,d1,d2,d3) and plotted separately for control and Samp1-KD samples as trajectory plots (mean ± SEM) using ggplot2 (v4.0.3) (Wickham, 2016).

### Gene set enrichment analysis

Gene set enrichment analysis (GSEA) was performed using fgsea (v1.32.4) (Korotkevich et al., 2021) on the ranked gene list from the KD vs. control comparison at day 3. Genes were ranked by the metric sign(log₂FC) × -log₁₀(p-value), capturing both direction and statistical effect. Gene sets for GO Biological Process (GO:BP) were retrieved from the Molecular Signatures Database (MSigDB) via the msigdbr package (v26.1) using Ensembl gene identifiers. Gene sets with fewer than 15 or more than 500 genes were excluded. Terms with adjusted p-value < 0.05 were considered significant.

### Integration with published DamID data

To identify genes that are transcriptionally repressed during normal myogenic differentiation and show increased association with the nuclear periphery, RNA-seq results were integrated with published Lamin B1 DamID data from Robson et al. (2016). Genes significantly downregulated in control cells from day 0 to day 3 (padj < 0.05, log₂FC < -1.5) that did not meet the same downregulation criteria in Samp1-KD cells (padj ≥ 0.05 or log₂FC ≥ -1.5) were matched to the published DamID dataset. Genes showing a positive gene-body Δlog₂ value (MT − MB DamID), indicating increased association with the nuclear periphery during differentiation, were visualized in the scatter plot.

### Statistical analysis

Statistical analyses and data visualization of microscopy and immunoblotting data were performed using GraphPad Prism version 11 (GraphPad Software). The biological replicate was used as the unit of statistical analysis, and unless otherwise indicated in the corresponding figure legend, data are presented as mean ± standard deviation (SD), with individual biological replicate values shown. Comparisons between replicate-matched conditions were performed using paired two-tailed t-tests, with each pair representing measurements obtained from the same biological experiment. Pairwise comparisons were planned according to the experimental design and specific biological questions, with only a limited number of comparisons performed for each outcome; therefore, no adjustment for multiple comparisons was applied. For immunoblotting, statistical analyses were performed on band intensities normalized to the corresponding β-actin loading control.

Statistical significance was defined as P < 0.05. Exact P values and the numbers of independent biological replicates are reported in the corresponding figure legends.

## Materials availability

The stable C2C12 cell lines and pTandemH reporter plasmid used in this study are available from the corresponding author upon reasonable request.

## Use of generative AI

ChatGPT (OpenAI) was used during manuscript preparation to assist with language editing, manuscript organization, and development and troubleshooting of custom image-analysis code. All code, analyses, scientific interpretations and final wording were reviewed and validated by the authors.

## Data Availability

The RNA-sequencing data generated in this study have been deposited in ArrayExpress (BioStudies), under accession number E-MTAB-17698. The dataset is currently under embargo and will be made publicly available upon publication. The published Lamin B1 DamID data analyzed in this study are available through GEO accession GSE80330, with the C2C12 Lamin B1 DamID dataset under GSE80328. Quantitative source data underlying the figures and original western blot images will be provided as source-data files with the manuscript.

## Code Availability

Custom code used for image analysis, DNA-FISH analysis, and FRIC analysis in this study will be available as part of the publication process.

## Funding

This work was supported by grants from Vetenskapsrådet (2020-2026), Olav Thons Stiftelse and Cancerfonden (2025-2027).

## Author Contributions

U.K., E. Hallberg: conceptualization and study design. E. Hallberg: supervision. E. Hedlund: supervision of RNA-seq and bioinformatic analyses. E. Hedlund, E. Hallberg: resources and funding acquisition. U.K.: cell culture and sample preparation, microscopy experiments, FRIC, DNA-FISH, quantitative image analysis, immunoblotting analysis, data curation, biological interpretation, final figure preparation, and writing of the original draft. I.M.: RNA-seq data processing and bioinformatic analysis, including differential expression analysis, module-score analysis, gene set enrichment analysis, generation of RNA-seq visualizations, and integration of the RNA-seq data with published Lamin B1 DamID data. S.G.A.: RNA-seq library preparation. S.A.: cell lysate preparation and western blotting. All authors contributed to editing and reviewing the manuscript and approved the final version.

## Competing Interests

The authors declare no competing interests.

## Figure Legends

Figure 1–figure supplement 1.

Control and Samp1-knockdown (KD) C2C12 cells were analyzed before differentiation induction (D0) and after 3 days of differentiation (D3). **(a)** Representative western blot and quantification of MyHC protein abundance. MyHC band intensities were normalized to the corresponding β-actin loading-control intensities. Three independent biological experiments were analyzed. **(b)** Representative western blot and quantification of Samp1 protein abundance normalized to β-actin. Four independent biological experiments were analyzed. Data are presented as mean ± SD, with individual symbols representing biological replicates. Statistical significance was assessed using paired two-tailed *t*-tests on replicate-matched values. Exact *P* values were as follows: **a**, control D0 versus KD D0, P = 0.3269; control D0 versus control D3, P = 0.0333; KD D0 versus KD D3, P = 0.3326; control D3 versus KD D3, *P* = 0.0328. **b**, control D0 versus KD D0, P = 0.0299; control D0 versus control D3, P = 0.0105; KD D0 versus KD D3, P = 0.3910; control D3 versus KD D3, P = 0.0019. *P < 0.05; **P < 0.01; ns, not significant.

Figure 3–figure supplement 1.

**(a)** Complete radial H2B/H3.3 heterochromatin-ratio profiles across all 40 concentric nuclear zones, extending from the nuclear periphery to the interior. FRIC analysis was performed in three independent biological experiments, with at least 14 nuclei analyzed per condition in each experiment. Lines show the mean H2B/H3.3 ratio, and error bars indicate SEM. **(b)** Chromosome 8 signal within the combined peripheral zones 1 + 2 and central zones 4 + 5, expressed as a percentage of total nuclear chromosome 8 fluorescence. Three independent biological experiments were analyzed, with at least 120 nuclei evaluated per condition in each experiment. Data are presented as mean ± SD, with individual symbols representing biological replicates. Statistical significance was assessed using paired two-tailed t-tests on replicate-matched values. Exact P values were as follows: peripheral zones 1 + 2: control D0 versus control D3, P = 0.0209; KD D0 versus KD D3, P = 0.9635; control D3 versus KD D3, P = 0.0376; central zones 4 + 5: control D0 versus control D3, P = 0.0075; KD D0 versus KD D3, P = 0.9967; control D3 versus KD D3, P = 0.0329. *P < 0.05; **P < 0.01; ns, not significant.

## Notes

### Competing Interest Statement

The authors have declared no competing interest.

### Summary of Updates

Accession number for ArrayExpress was added; Figure 1-Supplement 1 and Figure 3-Supplement 1 were added; figure legend for Figure 3-Supplement 1 was modified.

