## Supplementary figures and images for "The nuclear membrane protein Samp1 links peripheral genome organization to the myogenic transcriptional program"

### Figure 1 - Supplement 1

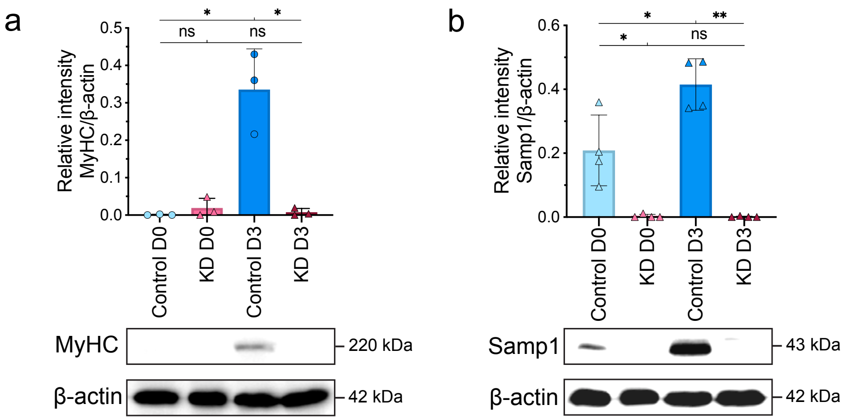

### Figure 3 - Supplement 1

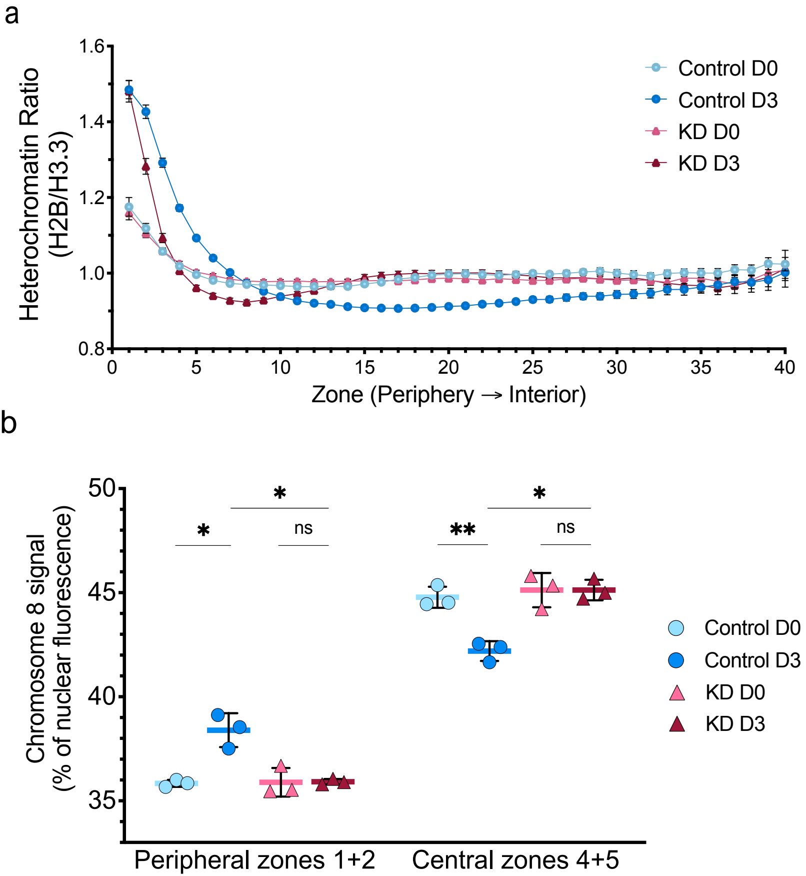
